# Metabolic Engineering of Oleaginous Yeast *Rhodotorula toruloides* for Overproduction of Triacetic Acid Lactone

**DOI:** 10.1101/2022.02.24.481788

**Authors:** Mingfeng Cao, Vinh G. Tran, Jiansong Qin, Andrew Olson, J. Carl Schultz, Chunshuai Huang, Dongming Xie, Huimin Zhao

## Abstract

The plant-sourced polyketide triacetic acid lactone (TAL) has been recognized as a promising platform chemical for the biorefinery industry. However, its practical application was rather limited due to low natural abundance and inefficient cell factories for biosynthesis. Here we report the metabolic engineering of oleaginous yeast *Rhodotorula toruloides* for TAL overproduction. We first introduced a 2-pyrone synthase gene from *Gerbera hybrida* (*GhPS*) into *R. toruloides* and investigated the effects of different carbon sources on TAL production. We then systematically employed a variety of metabolic engineering strategies to increase the flux of acetyl-CoA by enhancing its biosynthetic pathways and disrupting its competing pathways. We found that overexpression of citrate lyase (ACL1) improved TAL production by 45% compared to the *GhPS* overexpressing strain, and additional overexpression of acetyl-CoA carboxylase (ACC1) further increased TAL production by 29%. Finally, we characterized the resulting strain I12-*ACL1-ACC1* using fed-batch bioreactor fermentation in glucose or oilcane juice medium with acetate supplementation and achieved a titer of 28 g/L or 23 g/L TAL, respectively. This study demonstrates that *R. toruloides* is a promising host for production of TAL and other acetyl-CoA-derived polyketides from low-cost carbon sources.

**Graphical abstract:** 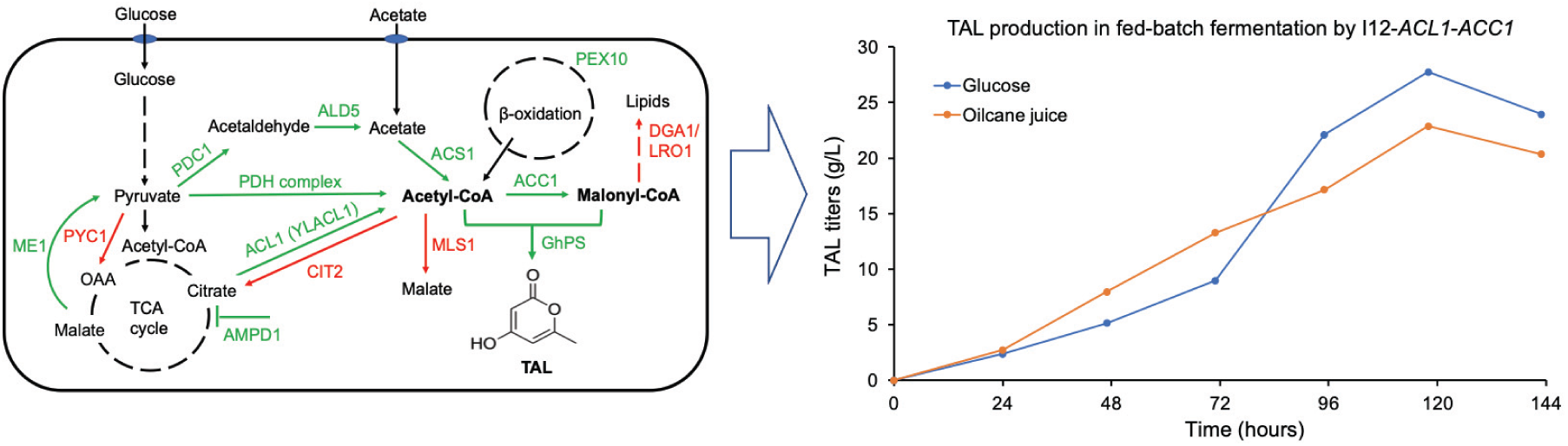

Triacetic acid lactone (TAL) is a promising platform chemical. Cao et al. overexpressed 2-pyrone synthase in oleaginous yeast *Rhodotorula toruloides* to produce TAL. They systematically evaluated various metabolic gene targets to increase acetyl-CoA and malonyl-CoA levels for TAL production and found that overexpression of both *ACL1* and *ACC1* led to 28 g/L or 23 g/L of TAL from glucose or oilcane juice with acetate supplementation, respectively, in fed-batch fermentation.

## 1. Introduction

The demand for renewable biobased products has attracted great interest in using microbial cell factories to replace traditional petroleum-based chemical manufacturing processes (Cardenas & Da Silva, 2014; Du et al., 2011; H. Liu et al., 2019; Markham et al., 2018). Among the chemicals that can be produced by microbial cell factories, polyketides represent a large family of naturally occurring metabolites with diversified industrial and biomedical applications. Due to the low yield by native producers and the challenge of introducing chiral centers by chemical catalysis, microbial bioconversion is considered a feasible approach for large-scale production of polyketides from renewable feedstocks (Keatinge-Clay, 2016; H. Liu et al., 2019; Robinson, 1991).

Triacetic acid lactone (TAL), also known as 4-hydroxy-6-methyl-2-pyrone, is a simple, yet interesting, polyketide. It has been demonstrated as a potential platform chemical for the production of commercially valuable bifunctional chemical intermediates and end products, including phloroglucinol (Zha et al., 2004), acetylacetone (Saunders et al., 2015), and sorbic acid (Chia et al., 2012). TAL is currently produced by chemical catalysis starting with the pyrolysis of acetic acid (Saunders et al., 2015). However, its industrial application was hampered by the detrimental catalysts and toxic byproducts (H. Liu et al., 2019). Therefore, it is highly desirable to develop an environment-friendly biological route for converting inexpensive substrates to TAL.

The biosynthesis of TAL is catalyzed by a type III polyketide synthase, 2-pyrone synthase (2-PS) via two iterative decarboxylation/condensation reactions using a starter acetyl-CoA and two extender malonyl-CoA molecules. 2-PS encoded by *g2ps1* gene was first isolated from *Gerbera hybrida* (Abe et al., 2005; Eckermann et al., 1998), and has been genetically introduced to conventional organisms *Escherichia coli* (Li et al., 2018; Tang et al., 2013; Xie et al., 2006) and *Saccharomyces cerevisiae* (Cardenas & Da Silva, 2014, 2016; Saunders et al., 2015; Sun et al., 2021) for TAL production. However, low titers were obtained in these conventional organisms due to cellular toxicity, limited intracellular acetyl-CoA pool, or unbalanced energy/cofactor supply (H. Liu et al., 2019).

Recent studies showed that oleaginous, nonconventional yeasts such as *Yarrowia lipolytica* and *Rhodotorula toruloides* (also known as *Rhodosporidium toruloides*) can be used for efficient TAL production due to their potential high flux through the key polyketide precursors, acetyl-CoA, and malonyl-CoA (Abdel-Mawgoud et al., 2018; Park et al., 2018). As a well-known lipid producer, *Y. lipolytica* was chosen for TAL biosynthesis via heterologous expression of 2-pyrone synthase (Yu et al., 2018), and the best engineered *Y. lipolytica* strain achieved a titer of 35.9 g/L TAL in 280 hours and a yield of up to 43% of the theoretical yield from glucose (Markham et al., 2018). *R. toruloides* is an oleaginous basidiomycete yeast, which can grow on various sugars and produce a broad range of lipid and nonlipid chemicals (Jagtap & Rao, 2018; Zhang et al., 2016; Zhang et al., 2021). Compared with *Y. lipolytica, R. toruloides* has a greater substrate range and natively produces TAG at much higher titers (Jagtap & Rao, 2018; Zhang et al., 2016; Zhang et al., 2021), but was less explored as a result of the unannotated genome sequence (Coradetti et al., 2018; Zhu et al., 2012) and lack of sophisticated genetic tools (Park et al., 2018). Nevertheless, the recent progress on characterization of constitutive promoters (Nora et al., 2019; Wang et al., 2016), development of CRISPR based genome editing tools (Jiao et al., 2019; Otoupal et al., 2019; Schultz et al., 2019), RNA interference tool (X. Liu et al., 2019), genome-scale model (Dinh et al., 2019), and functional genomics (Coradetti et al., 2018) enable us to perform metabolic engineering of *R. toruloides* for production of value-added compounds (Wen et al., 2020), specifically TAL.

In this study, we first expressed codon-optimized 2-PS genes from various organisms in *R. toruloides* and investigated the production of TAL under different culture conditions. We then created and characterized a broad set of TAL-producing overexpression and knockout gene targets in *R. toruloides* IFO0880. After combinatorial optimization of various targets, our final strain of I12-*ACL1*-*ACC1* achieved a maximum titer of 28 g/L within 120 hours in fed-batch fermentation from glucose with acetate addition. Then, we demonstrated the feasibility of olicane juice, an inexpensive carbon source as the substrate for TAL production, which produced 23 g/L TAL in fed-batch fermentation. This work not only establishes *R. toruloides* as a novel host organism for TAL biosynthesis but also demonstrates its potential as a biotechnological chassis for production of high-value chemicals from low-cost substrates.

## 2. Materials and methods

### 2.1 Strains, media, and chemicals

All strains used in this study are listed in Table 1. *E. coli* DH5α (New England Biolabs, Ipswich, MA) was used to maintain and amplify plasmids, and cells were grown in Luria Broth (LB) medium at 37 °C, 250 rpm with 100 µg/mL ampicillin or 50 µg/mL kanamycin. *R. toruloides* IFO0880 and its mutants were grown at 30 °C, 250 rpm in YPD media (1% yeast extract, 2% peptone, 2% glucose) for routine handling. For selection or maintenance of transformants, 200 µg/mL G418 (KSE Scientific, Durham, NC), 50 µg/mL hygromycin (InvivoGen, San Diego, CA) or 100 µg/mL nourseothricin (Gold Biotechnology, St. Louis, MO) was supplemented as necessary. TAL production media include YPD, YP2D (1% yeast extract, 2% peptone, 4% glucose), YPX (YP plus 2% xylose), YPDX (YP plus 1.4% glucose, 0.6% xylose), YPG (YP plus 2% glycerol), YPS (YP plus 2% sucrose), YP-NaAc (YP plus 2% sodium acetate) and SC (synthetic complete medium: 1.7 g/L yeast nitrogen base, 5 g/L ammonium sulfate, 0.78 g/L complete synthetic mixture, 2% glucose).

**Table 1.** List of plasmids and strains.

| Plasmids | Description | Source |
| --- | --- | --- |
| pRS426 | Used for cloning of pMB1 origin, *Amp*<sup>R</sup>, and *S. cerevisiae* elements of 2μ origin and *URA3* | Laboratory stock |
| pGI2 | *Kan*<sup>R</sup> for bacteria, *Nat*<sup>R</sup> for yeast, binary plasmid | (Abbott et al., 2013) |
| pGI2_880_ACC | Used for cloning of *ACC1* gene from *R. toruloides* | (Zhang et al., 2016) |
| pGI2-TEF1-X | Expression vector in *R. toruloides*, containing *Nat*<sup>R</sup>, *pTEF1* promoter and *T35S* terminator | This study |
| pRTN | Expression vector in *R. toruloides*, containing *Nat*<sup>R</sup>, *pANT* promoter and *T35S* terminator | Laboratory stock |
| pRTG2-X | Derived from pRTN, containing *G418*<sup>R</sup>, *pANT* promoter and *T35S* terminator | This study |
| pRTHyg-X | Derived from pRTN, containing *Hyg*<sup>R</sup>, *pANT* promoter and *T35S* terminator | This study |
| pRTH | gRNA expression vector in *R. toruloides*, containing *Hyg*<sup>R</sup>, *5S-tRNA*<sup>Tyr</sup>, 2 *BsaI* sites, and *SUP4* terminator for SgRNA | Laboratory stock |
| pRTH-X | Derived from pRTH with target N20 insertion at *BsaI* sites (no *BsaI* left) | This study |
| pRT-Cas9-SgRNA-Hyg | Derived from pRTH-X with *Cas9* expression cassette | This study |
| Strains | Description | Source |
| DH5α | Cloning host | NEB |
| IFO0880 | *R. toruloides* strain IFO0880, mating type A2 | NBRC, Japan |
| I12 | IFO0880 with genome integrated codon optimized *GhPS* cassette | This study |
| I12-*Cas9* | I12 with genome integrated codon optimized *Cas9* cassette | This study |
| I12-*ACL1* | I12 with genome integrated *ACL1* cassette | This study |
| I12-*ACL1-ACC1* | I12-*ACL1* with genome integrated *ACC1* cassette | This study |

LB broth, bacteriological grade agar, yeast extract, peptone, yeast nitrogen base, ammonium sulfate, and D-xylose were obtained from Difco (BD, Sparks, MD), while complete synthetic medium was purchased from MP Biomedicals (Solon, OH). TAL standard was purchased from Aldrich Chemical Co. (Milwaukee, WI). All restriction endonucleases, Q5 DNA polymerase, and Gibson Assembly Cloning Kit were purchased from New England Biolabs (Ipswich, MA). The QIAprep spin mini-prep kit was purchased from Qiagen (Valencia, CA), the Wizard Genomic DNA Purification Kit was purchased from Promega (Madison, WI), whereas Zymoclean Gel DNA Recovery Kit and Zymoprep Yeast Plasmid Miniprep Kits were purchased from Zymo Research (Irvine, CA). All other chemicals and consumables were purchased from Sigma (St. Louis, MO), VWR (Radnor, PA), and Fisher Scientific (Pittsburgh, PA). Sequences for key primers, N20 of SgRNA, and gene targets (Coradetti et al., 2018) were summarized in Table S1 and Table S2. Primers were synthesized by Integrated DNA Technologies (IDT, Coralville, IA), while heterologous genes were codon optimized by GeneOptimizer or the JGI BOOST tool and synthesized by GeneArt (Invitrogen, CA) or Twist Bioscience (San Francisco, CA). gRNAs were designed using the CRISPRdirect (https://crispr.dbcls.jp/) or the Benchling gRNA tool. DNA sequencing was performed by ACGT, Inc. (Wheeling, IL). Plasmid mapping and sequencing alignments were carried out using SnapGene software (GSL Biotech, available at snapgene.com).

### 2.2 Plasmid construction

#### Plasmids for 2-PS expression

The codon-optimized 2-PS genes were synthesized with two homologous ends to *Mfe*I and *Spe*I digested pGI2 (Abbott et al., 2013; Zhang et al., 2016) backbone, which contains nourseothricin resistance (*NAT*^R^) for yeast, and then assembled with *pTEF1* promoter and *T35S* terminator for gene expression by Gibson assembly in *E. coli* (Gibson et al., 2009).

#### Plasmids for overexpression of metabolic gene targets

The plasmid pRTG2-X (X represents gene expression targets) for gene targets expression was constructed based on a previously developed plasmid pRTN, which contains the *E. coli* genetic elements of pUC19 (pMB1 origin, ampicillin resistance), the *S. cerevisiae* genetic elements of pRS426 (2µ origin and *URA3* selection marker), the strong *R. toruloides p17* or *pANT* promoter, the target gene, the *T35S* terminator, and a *R. toruloides* G418 resistance (*G418*^R^) cassette from NM9 (Schultz et al., 2019) using DNA assembler (Shao et al., 2009). The multiple gene expression plasmid pRTHyg-X was pieced together from NM8 (Schultz et al., 2019) for hygromycin resistance (*Hyg*^R^), *pANT* promoter and *Tncbt* for gene expression.

#### Plasmids for gene target knockout

The previously constructed plasmid pRTH-X (X represents gene knockout targets) was used for gRNA cloning and expression, which contains the *E. coli* genetic elements (pMB1 origin, ampicillin resistance), the *S. cerevisiae* genetic elements of pRS426 (2µ origin and *URA3*), a gRNA expression cassette with the IFO0880 5S rRNA, tRNA^Tyr^, N20 (targeting the first 10% ORF), the *S. cerevisiae* SUP4 terminator, and a *R. toruloides Hyg*^R^ cassette from pZPK-PGPD-HYG-Tnos (Lin et al., 2014). For multiple gene knockout, the plasmid pRT-Cas9-SgRNA-Hyg was constructed from pRTH-X with an integrated *Cas9* expression cassette.

### 2.3 Yeast transformation

Most of the linear fragments for *R. toruloides* transformation were generated by PCR amplification of the genes or gRNA expression cassettes, together with the selection markers using the primers ZPK F/R, or gRNA F/R (Table S1), respectively, whereas the fragments with sizes larger than 8 kb (e.g., *ACC1-G418/Hyg*) were excised from the plasmids by restriction enzyme digestion. Fragments were then cleaned using DNA Clean & Concentrator-5 Kit (Zymo Research) before transforming to *R. toruloides*.

*R. toruloides* was transformed using heat shock as previously described (Otoupal et al., 2019; Schultz et al., 2021). Briefly, a single colony was picked and cultured overnight at 30 °C in 3 mL YPD medium supplemented with an appropriate antibiotic if required. The overnight culture was transferred to 50 mL fresh YPD with an OD_600_ of 0.2 and cultured for another 4 hours at 30 °C to an OD_600_ of approximately 1.0. Cells were collected by centrifugation, washed twice with sterile water and once with 100 mM LiAc (pH 7.6) (Sigma Aldrich, St. Louis, MO), and then resuspended in a transformation mixture of 240 µL PEG3350 (Sigma Aldrich, St. Louis, MO), 36 µL 1 M lithium acetate, 50 µL of 2 mg/mL salmon sperm DNA (Sigma Aldrich, St. Louis, MO), and 1-2 µg of linear DNA dissolved in 34 µL of water. The cells were incubated with 200 rpm shaking in the mixture for 30 min at 30 °C. Then, 34 µL of dimethyl sulfoxide (DMSO) was added to the mixture, which was briefly vortexed, and heat shocked at 42 °C for 20 min. The cells were pelleted, washed with YPD, resuspended in 2 mL YPD, and recovered overnight with shaking. Cells were then collected and plated on YPD solid medium supplemented with the appropriate antibiotic(s). To verify the integrated fragments, genomic DNA was extracted using the Wizard Genomic DNA Purification Kit (Promega), and the target locus was PCR amplified for sequencing.

### 2.4 Culture tube or shake flask fermentation

Single colonies were picked and cultured for 24 hours at 30 °C in 3 mL YPD liquid medium supplemented with appropriate antibiotics in 14 mL culture tubes (VWR) as seed culture. Fermentations were inoculated from seed culture to media with alternative carbon sources at an initial OD_600_ of 0.2 and grown for an additional 72 hours prior to sample preparation. For acetate spike, filter sterilized 20x sodium acetate (NaAc) was added to media at 12 h. Samples were collected by centrifuge and diluted 20 times for high performance liquid chromatography (HPLC) analysis. Since a deficient cell growth and tiny amounts of TAL were obtained in YP (1% yeast extract and 2% peptone) medium, yield was calculated based on the produced TAL over all carbons of sugars and/or acetate.

### 2.5 Fed-batch fermentation

For fed-batch fermentation in bioreactors, single colonies of *R. toruloides* I12-*ACL1* and I12-*ACL1*-*ACC1* were used to inoculate shake flask cultures with each containing 50 mL of YPAD medium supplemented with appropriate antibiotics, as described in section 2.1. Cells in the flask cultures were grown at 30 °C and 250 rpm until the OD_600_ reached 2-5. The seed culture from each flask was transferred to a 1-L bioreactor (Biostat B-DCU, Sartorius, Germany) with 0.7 L fermentation medium, which contained 15 g/L yeast extract, 15 g/L peptone, 10 g/L (NH_4_)_2_SO_4_, 6 g/L KH_2_PO_4_, 2 g/L Na_2_HPO_4_, 1 mL/L Antifoam 204 (Sigma-Aldrich), 1.5 mg/L Thiamin·HCl, 1.2 g/L MgSO_4_, 2 mL/L trace metals (100X), 50 g/L glucose, and appropriate antibiotics (same as described in section 2.1). The trace metal (100X) solution contained 10 g/L citric acid, 1.5 g/L CaCl_2_·2H_2_O. 10 g/L FeSO_4_·7H_2_O, 0.39 g/L ZnSO_4_·7H_2_O, 0.38 g/L CuSO_4_·5H_2_O, 0.2 g/L CoCl_2_·6H_2_O, and 0.3 g/L MnCl_2_·4H_2_O. The dissolved oxygen level was maintained at 20% of air saturation, and the temperature was set at 30 °C. The pH value was controlled at 6.0 using glacial acetic acid and 10 M KOH. When the residual glucose concentration decreased to nearly 0 g/L (as indicated by a sharp decreasing in agitation speed and an increase in pH value), continuous feeding of glucose (from a 600 g/L stock solution) was used to maintain its residual concentrations within 20 g/L. Sodium acetate (460 g/L) was fed to the fermenter in pulse at different time points: 10 mL at 36, 48, 60, 72, and 96 hours, and 8 mL at 108, 120, 132, 144, 156, and 168 hours, respectively. The fed-batch fermentation was conducted in biological duplicates.

For the fed-batch fermentation experiments with oilcane, the original oilcane feed solution containing a total sugar of 152 g/L (68.9 g/L glucose, 61.6 g/L fructose, and 21.5 g/L sucrose) was concentrated to about 450 g/L total sugar by evaporation through the boiling at atmospheric pressure. The concentrated oilcane feed solution was further autoclaved at 121 °C for 30 min before it was used to provide initial sugars in the medium and to feed sugars during the fed-batch fermentation. The initial medium contained a total oilcane sugar of 50 g/L and all other medium components, as described previously for the glucose fed-batch fermentation. The oilcane feeding started when the initial sugars were depleted, as indicated by a sharp decrease in agitation speed and an increase in pH value. Oilcane was fed to control the residual glucose concentrations within 0∼10 g/L. All other fermentation conditions, including acetate feeding, were same as previously described for the fed-batch fermentation experiments with glucose.

### 2.6 Analytical methods

Samples were prepared by diluting in methanol to the linear range, vortex mixing, and centrifuging at 16,000 *g* for 5 min to remove cells. After being filtered by a 0.2 μm filter, the supernatant was injected into the HPLC for TAL, sugars, and acetate analyses. (1) *TAL characterization*. The analytical HPLC was carried out on an Agilent 1260 Infinity series instrument equipped with a diode array detector (DAD) using a Phenomenex Kinetex^®^ 5 µm EVO C18 100 Å LC column (150 × 4.6 mm; Phenomenex, USA). The solvent system comprises solvent A (water supplemented with 0.1% trifluoroacetic acid) and B (acetonitrile supplemented with 0.1% trifluoroacetic acid). The elution process runs the following program: 2% B to 7% B (linear gradient, 0–5 min), 7% B to 95% B (5–6 min), 95% B (isocratic elution, 6–8 min), 95% B to 2% B (8–9 min), 2% B (isocratic elution, 9–11 min). Full wavelength scanning (UV/Vis) and Liquid Chromatography-Mass Spectrometry (LC-MS) were performed to determine the specific absorbance and molecular weight of the target products using >98.0% purity TAL as a reference. LC-MS analysis was running on a Waters Synapt G2-Si ESI/LC-MS (Milford, MA), equipped with ESI positive ion mode (Bruker, Amazon SL Ion Trap) and a Kinetex 2.6-μm XB-C18 100 Å (Phenomenex). (2) *Sugar and acetate analyses*. Glucose, xylose, glycerol, sucrose, and acetate consumptions were measured using an Agilent 1260 Infinity HPLC (Santa Clara, CA), equipped with Rezex™ ROA-Organic Acid H^+^ (8%) column (Phenomenex Inc., Torrance, CA) and a refractive index detector (RID). The column and detector were run at 50 °C and 0.6 mL/min of 0.005 N H_2_SO_4_ was used as the mobile phase (J.-J. Liu et al., 2019).

## 3. Results and discussion

### 3.1 *R. toruloides* can serve as a TAL producer

High lipid production in oleaginous organisms like *R. toruloides* suggests a great potential for these organisms to synthesize alternative acetyl-CoA-derived products, specifically type III polyketide, TAL (Markham et al., 2018; Park et al., 2018; Wen et al., 2020). It was reported that *R. toruloides* could grow normally under harsh conditions (Lyu et al., 2021) or cultures with non-native products, including fatty alcohols (Liu et al., 2020), fatty acid ethyl esters (Zhang et al., 2021), and limonene (Liu et al., 2021). The product tolerance assay also showed that *R. toruloides* possessed a similar growth profile in YPD and YPD with 5 g/L TAL but longer lag and log phases in YPD with 7 g/L TAL supplementation (Fig. S1). Therefore, *R. toruloides* can be potentially engineered to produce high titers of TAL without significantly detrimental growth effects.

As the 2-pyrone synthase (2-PS) gene tested for TAL biosynthesis in yeast and bacteria was mainly from *Gerbera hybrida* (GhPS, Uniprot ID: P48391), we sought to explore additional 2-PS genes from a range of alternative organisms and characterize their function in *R. toruloides*. To assist the selection of 2-PS genes, the Enzyme Function Initiative-Enzyme Similarity Tool (EFI-EST) was used to create the sequence-similarity network (SSN) based on the InterPro family IPR011141 (Type-III polyketide synthase) (Gerlt et al., 2015). *GhPS* and three additional 2-PS genes from previously unexplored species, *Vitis vinifera* (Protein ID: CK203_022254), *Sphaceloma murrayae* (Protein ID: CAC42_3419), and *Aspergillus oryzae* (Protein ID: AO090701000566) were codon-optimized using the most frequently used codon (Table S2), synthesized, and cloned to pGI2 under the *pTEF1* promoter. The PCR-amplified *2-PS*-*NAT* cassettes were transformed and randomly genome-integrated *via* NHEJ to *R. toruloides*. However, of these genes, only *GhPS* produced TAL (Fig. S2). The strain I12 with one copy *GhPS* expression produced 2.0 ± 0.1 g/L TAL in YPD medium at 72 h in a culture tube, which was close to that obtained from the *Y. lipolytica* strain (2.1 g/L) with four copies of *GhPS* expression in a defined synthetic medium (Markham et al., 2018). Although codon optimization algorithms and genome integration loci may affect the *GhPS* expression, it indicates *R. toruloides* can serve as a promising platform for TAL production.

### 3.2 TAL production using various substrates

To evaluate the effects of substrates on TAL production, we chose the commonly used carbon sources, including xylose (X), glycerol (G), sucrose (S), complete synthetic medium (SC), and glucose/xylose mixed sugar (DX) (Fig. 1A). The results showed that YPD was a more preferred medium than SC, with a 4-fold higher TAL titer; compared to glucose, glycerol and glucose/xylose produced 5%∼10% higher TAL, while xylose and sucrose decreased TAL production. We also observed deficient cell growth and residual sugars in YPX, YPS, and SC media (Table S3), indicating a positive correlation between cell growth and TAL titer.

**Fig. 1.**
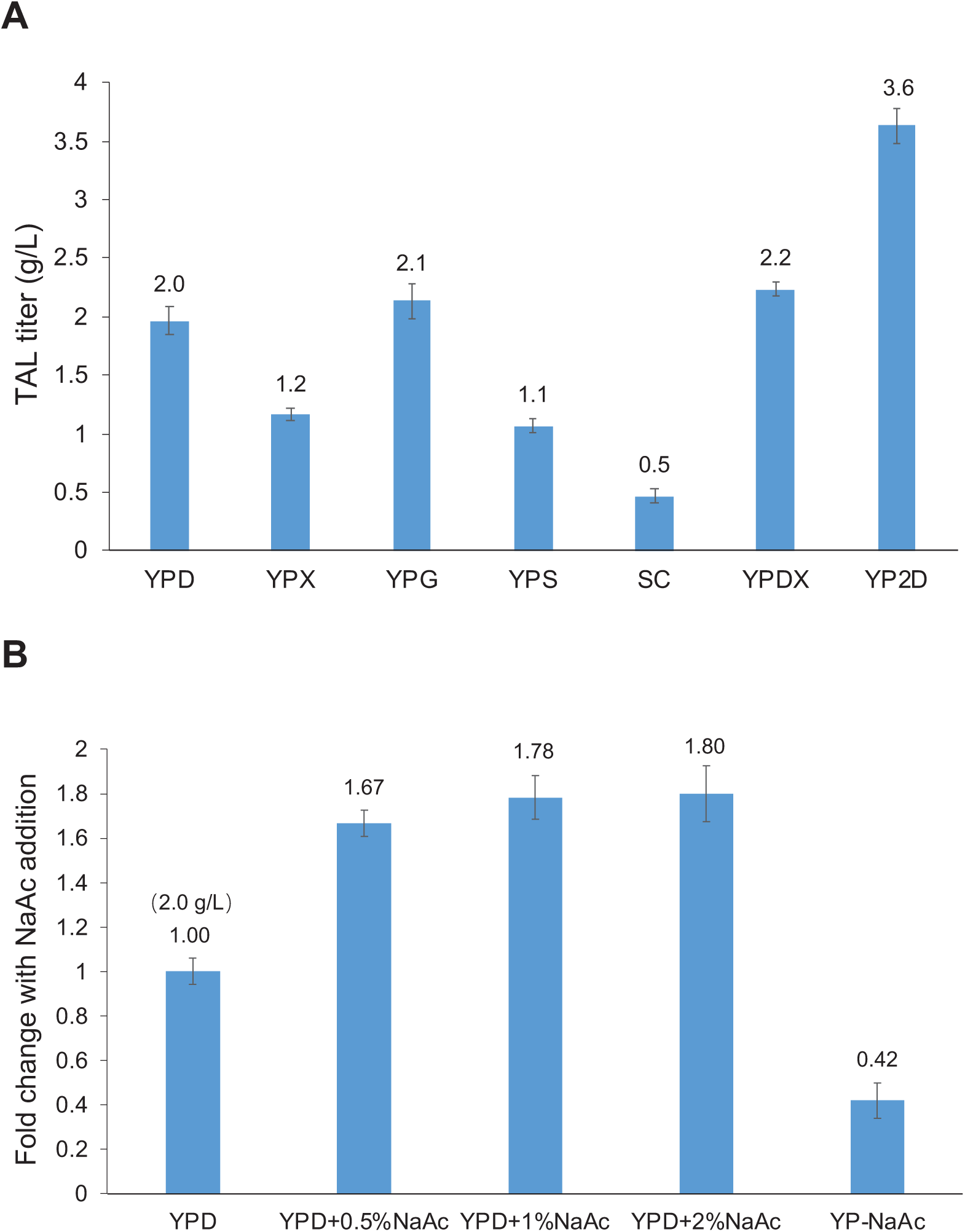
TAL production in *R. toruloides*-I12 using different substrates. (A) Commonly used sugars; (B) Acetate spiking affects TAL production.

It has been demonstrated that acetate feeding was beneficial to acetyl-CoA supply and TAL biosynthesis in *S. cerevisiae* and *Y. lipolytica* (H. Liu et al., 2019; Markham et al., 2018; Sun et al., 2021). We therefore supplemented 0.5%, 1%, and 2% NaAc to YPD with the 12 h *R. toruloides* cell culture and observed significant improvements, 67%∼80% higher TAL production under these spiking conditions, representing a similar titer to that of YP2D (3.6 g/L TAL from YP-4% glucose at 72 h) and ∼30% of the theoretical yield calculated from both glucose (2%) and acetate (0.5%) in culture tube (Fig. 1B).

To explore the potential role of acetate during TAL biosynthesis, we provided 2% NaAc as an alternative carbon source. However, the TAL produced from YP-NaAc was only 42% of that produced from YPD (Fig. 1B). The residual amounts of NaAc were also measured for the above-mentioned NaAc media, and only YPD+0.5%NaAc showed depletion of acetate after fermentation while a portion of acetate left in YPD+1%NaAc, YPD+2%NaAc, and YP-NaAc (Table S3). The acetate consumption indicated that acetate may not only act as a substrate for TAL production, but also be associated with the redox and regulatory mechanism, which has been elaborated in *Y. lipolytica* (Markham et al., 2018).

### 3.3 Single gene target engineering to improve TAL production

It is generally recognized that malonyl-CoA is the limiting precursor for polyketide synthase (Xu et al., 2011; Zha et al., 2009). Therefore, we overexpressed the endogenous acetyl-CoA-carboxylase (ACC1) to test whether the conversion of acetyl-CoA to malonyl-CoA would facilitate TAL synthesis (Fig. 2). As shown in Fig. 3A, overexpression of *ACC1* did not markedly enhance TAL production with only 6% improvement in YP2D at 120 h compared to that of the starting strain I12. To further drive the condensation of acetyl-CoA and malonyl-CoA, we introduced a second copy of *GhPS* gene via genome integration and achieved 4.8 g/L TAL at 120 h, ∼11% higher than that of I12 (Fig. 3A). Based on these results, we deduced that the limiting precursor for TAL overproduction was acetyl-CoA instead of malonyl-CoA. Therefore, we sought to increase the flux of acetyl-CoA by enhancing its biosynthetic pathways and disrupting its competing pathways.

**Fig. 2.**
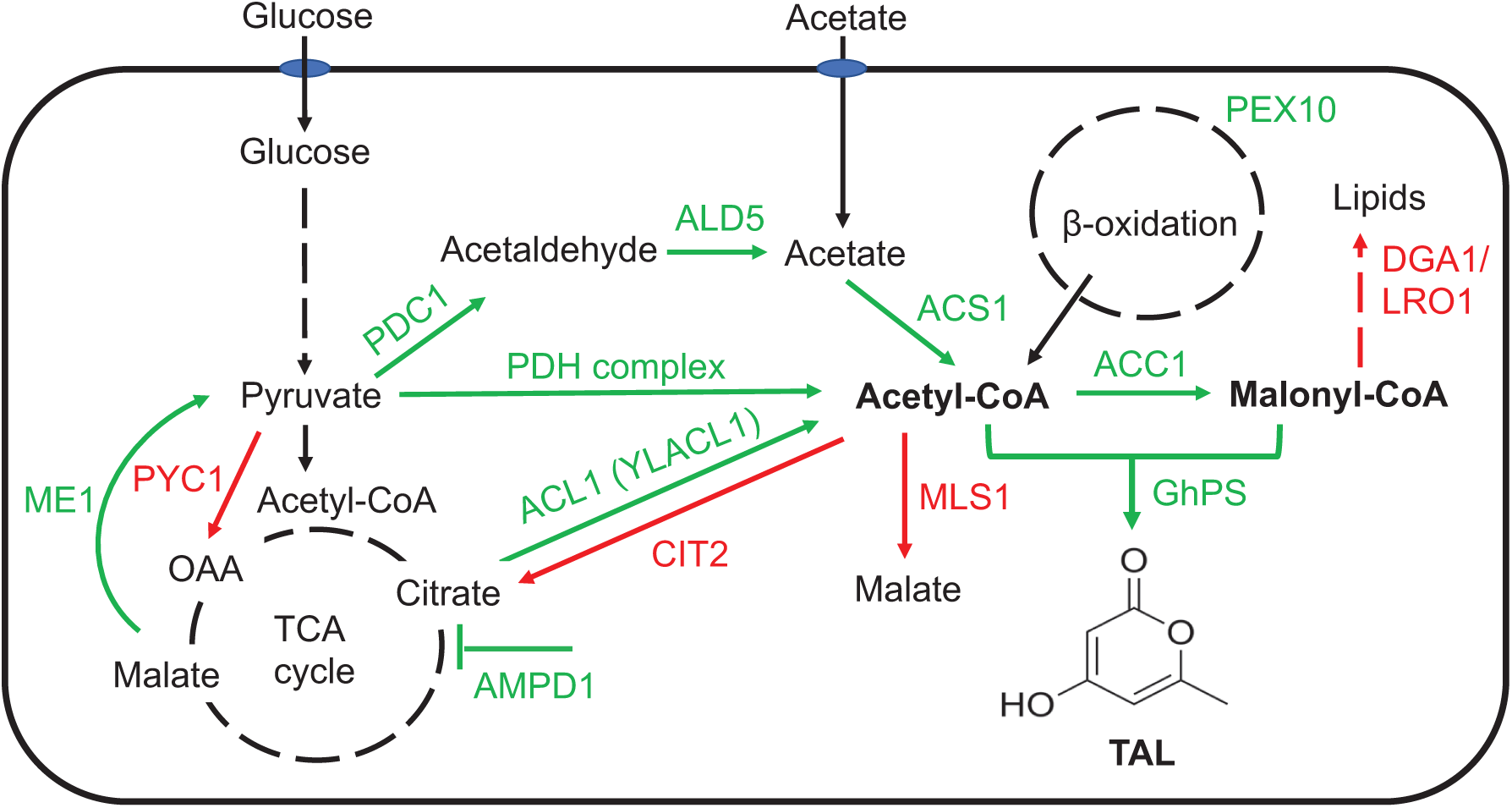
Metabolic pathway engineering for TAL biosynthesis in *R. toruloides*. PDC1, pyruvate decarboxylase; ALD5, acetylaldehyde dehydrogenase; ACS1, acetyl-CoA synthetase ACS1, ACC1, acetyl-CoA carboxylase; GhPS, 2-pyrone synthase gene from *Gerbera hybrida*; PDH, pyruvate dehydrogenase; ACL1, citrate lyase; YLACL1, citrate lyase from *Yarrowia lipolytica*; AMPD1, AMP deaminase; ME1, malic enzyme; PEX10, peroxisomal matrix protein; PYC1, pyruvate carboxylase; MLS1, cytosolic malate synthase; CIT2, peroxisomal citrate synthase; DGA1, diacylglycerol acyltransferase; LRO1, lecithin cholesterol acyltransferase. Note: Some metabolites were not positioned following their intracellular compartmentalization.

**Fig. 3.**
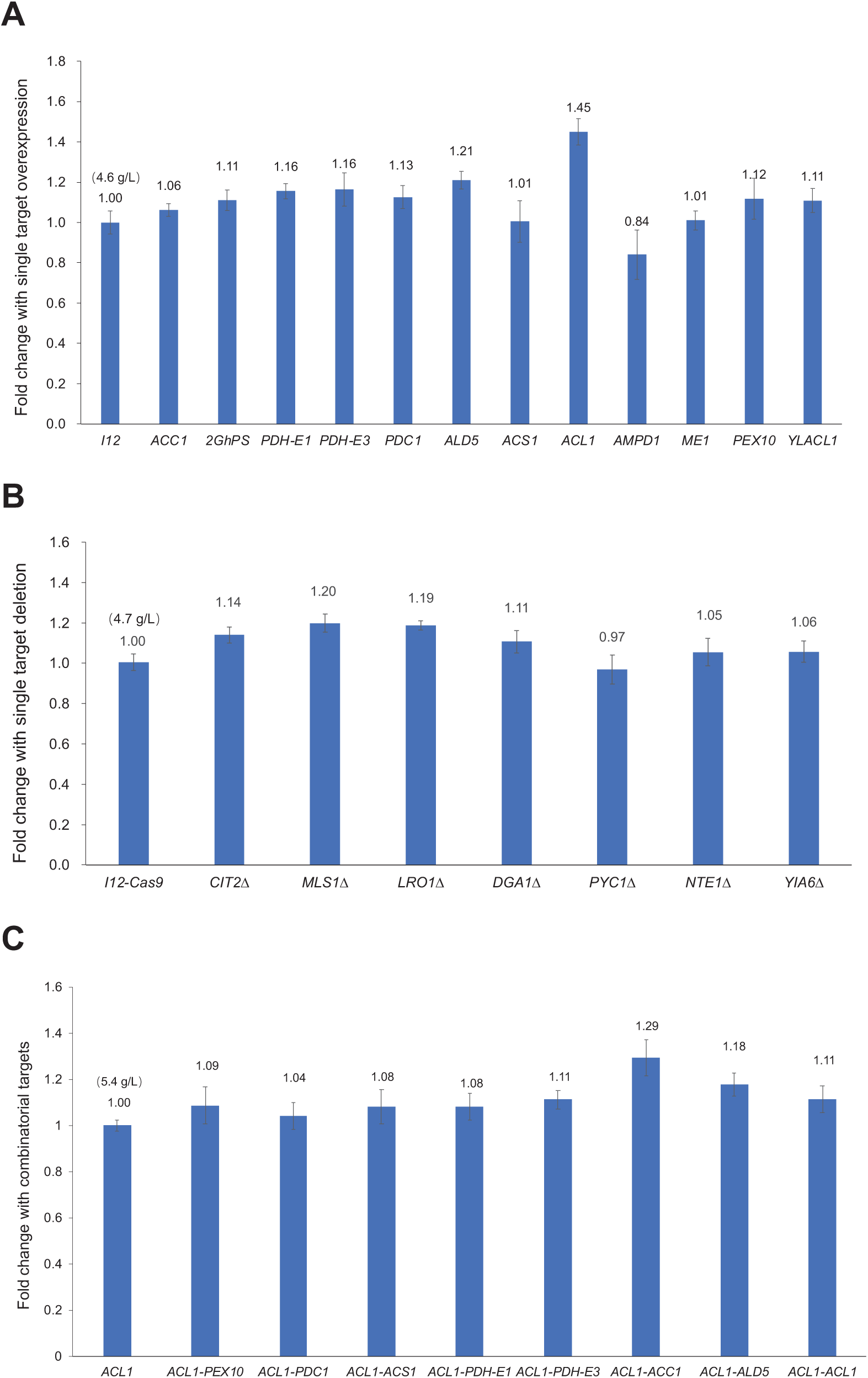
TAL production from different metabolic engineering strategies. A. Overexpression of selected gene targets; B. Disruption of selected gene targets. C. Multiple gene targets by combinatorial engineering.

#### 1) Enhancing acetyl-CoA biosynthetic pathways

We explored three distinct metabolic engineering strategies and characterized the roles of associated gene targets in TAL production (Fig. 3). First, we investigated the pyruvate dehydrogenase (PDH) complex pathway and overexpressed its subunits, E1 and E3 (*LPD1*) in I12 strain. Fermentation showed that both E1 and E3 overexpression improved TAL production by ∼16%, which reached ∼5.3 g/L at 120 h (Fig. 3A and Table S4). It is known that the PDH complex is located in the mitochondrial matrix in eukaryotes, and its compartmentalization is mediated via mitochondrial targeting sequence (MTS). However, the improved TAL production indicated that the tested subunits of E1 and E3 may not contain a fully featured MTS, or there may be leaky expression of these two subunits in the cytoplasm, which was similar to that of the overexpression of PDH complex in *S. cerevisiae* (Lian et al., 2014) and *Y. lipoytica* (Markham et al., 2018). As we failed to construct the mutants of other subunits of PDH, overexpression of the complete PDH or a cytoplastic PDH complex (i.e., *E. coli* cytoPDH) (Cardenas & Da Silva, 2016; Kozak et al., 2014) can be a potential strategy to increase the acetyl-CoA level.

Second, we evaluated the pyruvate dehydrogenase bypass pathway, which converts pyruvate to acetyl-CoA through a three-step reaction sequentially catalyzed by pyruvate decarboxylase (PDC), acetaldehyde dehydrogenase (ALD), and acetyl-CoA synthetase (ACS). The corresponding genes, *PDC1, ALD5*, and *ACS1* were individually overexpressed in I12 (Fig. 2). As shown in Fig. 3A and Table S4, overexpression of *PDC1* and *ALD5* improved TAL production by 13% and 21%, reaching 5.1 g/L and 5.5 g/L in YP2D at 120 h, respectively. However, no TAL improvement was achieved by *ACS1* overexpression, which is similar to the study in *S. cerevisiae* where *ACS1* overexpression did not improve *n*-butanol production because of low activity or post-translational deactivation (Lian et al., 2014).

Third, we explored the citrate route, a pathway that generates cytosolic acetyl-CoA from citrate and was reported to be present only in oleaginous yeasts (Pomraning et al., 2019; Vorapreeda et al., 2012; Zhu et al., 2012). The pathway gene *ACL1*, encoding ATP-citrate lyase has been overexpressed to increase lipid production in *Y. lipolytica* (Blazeck et al., 2014; Wang et al., 2015). A multi-omic analysis of *R. toruloides* also revealed that *ACL1* was expressed at extremely high level during lipogenesis stage (Zhu et al., 2012). Therefore, the endogenous *ACL1* was overexpressed in I12 strain (Fig. 2), and the TAL production was dramatically improved by 45%, to 6.6 g/L in YP2D in a test tube at 120 h (Fig. 3A and Table S4), which was ∼35% of the theoretical yield.

In addition, we overexpressed metabolic targets that could indirectly increase metabolic flux of acetyl-CoA, including *AMPD1* (encoding AMP deaminase) (Zhang et al., 2019), *ME1* (encoding malic enzyme), *PEX10* (encoding peroxisomal matrix protein), and *YLACL1* (ACL1 from *Y. lipolytica* and sequence included in Table S2) (Blazeck et al., 2014) (Fig. 2). The results showed that *PEX10* and *YLACL1* overexpression increased TAL titer by 12% and 11%, respectively, whereas *AMPD1* decreased TAL titer and *ME1* had no effect on TAL production (Fig. 3A). This suggests that up-regulation of β-oxidation by enhancing peroxisome biogenesis through *PEX10* overexpression is an alternative way to recycle acetyl-CoA for TAL production in *R. toruloides*.

#### 2) Disrupting acetyl-CoA competing pathways

Removing acetyl-CoA consuming pathways was demonstrated as an effective way to increase the availability of acetyl-CoA. In yeast, the glyoxylate shunt allows acetyl-CoA to be converted into a C4 carbon without carbon loss (Dolan & Welch, 2018). Therefore, we performed the inhibition of two key reactions of the glyoxylate cycle, namely peroxisomal citrate synthase, encoded by *CIT2*, and cytosolic malate synthase, encoded by *MLS1* (Chen et al., 2013), by a previously developed CRISPR/Cas9 method (Schultz et al., 2019). As shown in Fig. 3B, compared with I12-*Cas9* strain, the deletion of *CIT2* and *MLS1* improved TAL production by 14% and 20%, respectively.

In addition, we investigated the effects of disrupting other gene targets, including two acyltransferases (encoded by *DGA1*/*LRO1*), pyruvate carboxylase (encoded by *PYC1*), serine esterase or patatin-domain-containing protein (encoded by NTE1), and mitochondrial NAD^+^ transporter (encoded by *YIA6*). Among them, *DGA1* and *LRO1* are involved in TAG formation in *Y. lipolytica* (Athenstaedt, 2011), and *PYC1, NTE1*, and *YIA6* were reported to improve TAL production in *S. cerevisiae* (Cardenas & Da Silva, 2014). The fermentation showed that the deletion of *DGA1* and *LRO1* improved TAL titer by 11% and 19%, respectively, while the deletion of *NTE1, YIA6*, and *PYC1* had a marginal effect on TAL production (Fig. 3B), which is inconsistent with the observation in *S. cerevisiae*.

### 3.4 Multiple gene target engineering to improve TAL production

To further investigate the effects of multiple gene targets on TAL production in a combinatorial manner, we selected the top targets that improved TAL production more than 12%, i.e., *ACL1, ALD5, MLSΔ, LRO1Δ, PDH-E3, PDH-E1, CIT2Δ, PDC1*, and *PEX10* for the second round of metabolic engineering based on I12-*ACL1* strain. In addition, we included *ACC1* as its overexpression may result in improved malonyl-CoA concentration in an acetyl-CoA enhanced strain, I12-*ACL1*. We successfully obtained the mutant strains that overexpressed *ACL1, ALD5, PDH-E3, PDH-E1, PDC1, PEX10*, and *ACC1* through random genome integration using HYG selection, and the optimal combination was *ACL1*-*ACC1*, which produced 6.9 g/L TAL in YP2D at 120 h, representing a 29% improvement compared with I12-*ACL1* strain (Fig. 3C and Table S4). Unfortunately, we failed to obtain the correct mutants with *MLS1, LRO1, CIT2* deletion after transforming linear fragments containing *Cas9*-*SgRNA*-*Hyg* into I12-*ACL1* strain. Although a decent number of colonies were growing on YPD+HYG plates, which meant the *Hyg* expression cassette was integrated into genome, none of colonies had the expected genome mutation or *indel* after sequencing-based genotyping. Compared to the high genome editing (gene knockout) efficiency through transforming *SgRNA-Hyg* to I12-*Cas9* strain, the low editing efficiency in I12-*ACL1* may be caused by the co-expression of *Cas9* and *SgRNA* in a single fragment of *Cas9*-*SgRNA*-*Hyg*, which could result in the toxicity or lethality to the host cells. Therefore, a two-step transformation of individual *Cas9* and *SgRNA* could be beneficial for genome editing in I12-*ACL1*, and future efforts will be made to develop other antibiotic or auxotrophic selection markers in *R. toruloides*, or introduce recyclable expression platforms, such as a replicable episomal plasmid (Schultz et al., 2021) or a Cre/loxP site-specific recombination system (Díaz et al., 2018).

### 3.5 Scale-up TAL production in bioreactor

To evaluate the possibility of scaling up fermentation, fed-batch fermentation was performed using strains I12-*ACL1* and I12-*ACL1*-*ACC1*, which showed high TAL titers in culture tube fermentations. Before bioreactor fermentation, we performed shake flask cultures of I12-*ACL1* and I12-*ACL1*-*ACC1* in YP2D with 0.5% or 1% sodium acetate spike. The results showed that the cell OD_600_ and TAL production with 5 g/L sodium acetate addition were higher than that with 10 g/L sodium acetate (Fig. S3), and I12-*ACL1*-*ACC1* produced 7.1 g/L TAL at 96 h, which was 40% more than that of I12-*ACL1* in YP2D-0.5% NaAc. The yield of TAL production in YP2D-0.5% NaAc for I12-*ACL1*-*ACC1* reached 35% of the theoretical yield, which was similar to culture tube fermentation of I12-*ACL1*-*ACC1* in YP2D. However, it took 24 hours less fermentation time, indicating a higher oxygen concentration may benefit TAL production.

For bioreactor fermentation, *R. toruloides* strain I12-*ACL1*-*ACC1* was set up for scale-up using fed-batch cultures in medium containing yeast extract, peptone, glucose, and other trace metals, and produced 28 g/L of TAL from glucose-based medium at 118 h, representing a high volumetric productivity of 0.24 g/L/h (Fig. 4A and 4B, Table S5). The highest yield of 0.074 g TAL /g carbon source (glucose and acetate) for I12-*ACL1*-*ACC1* was also achieved at 118 h, which was 16% of the theoretical yield. Furthermore, to demonstrate the feasibility of fermentation using low-cost feedstocks and to take advantage of the capability of *R. toruloides* to assimilate a diverse range of substrates, including monosaccharides, oligosaccharides, and organic acids, we performed fed-batch fermentation using oilcane juice, a first-generation feedstock which is comprised of 152 g/L total sugars. As shown in Fig. 4C, I12-*ACL1*-*ACC1* strain grew well in the oilcane juice, suggesting they may have high tolerance towards the inhibitors present in the oilcane juice. The titer and yield of TAL were 23 g/L and 0.089 g/g carbon sources (glucose, fructose, sucrose, and acetate), representing 19% of the theoretical yield (Fig. 4D and Table S5), while the productivity was 0.19 g/L/h before 120 h. Although the yields in glucose-and oilcane juice-based fed-batch fermentation were relatively low, the volumetric productivities in both conditions were much higher than that in culture tube (0.058 g/L/h) or shake flask (0.074 g/L/h), and even higher than that reported in *Y. lipolytica* (0.12 g/L/h) (Markham et al., 2018). Meanwhile, the fed-batch fermentation provided us more knowledge of improving TAL production by optimizing fermentation conditions, including feeding rates, dissolved oxygen/pH control (Sun et al., 2021), and *in situ* product separation (Lee et al., 2016).

**Fig. 4.**
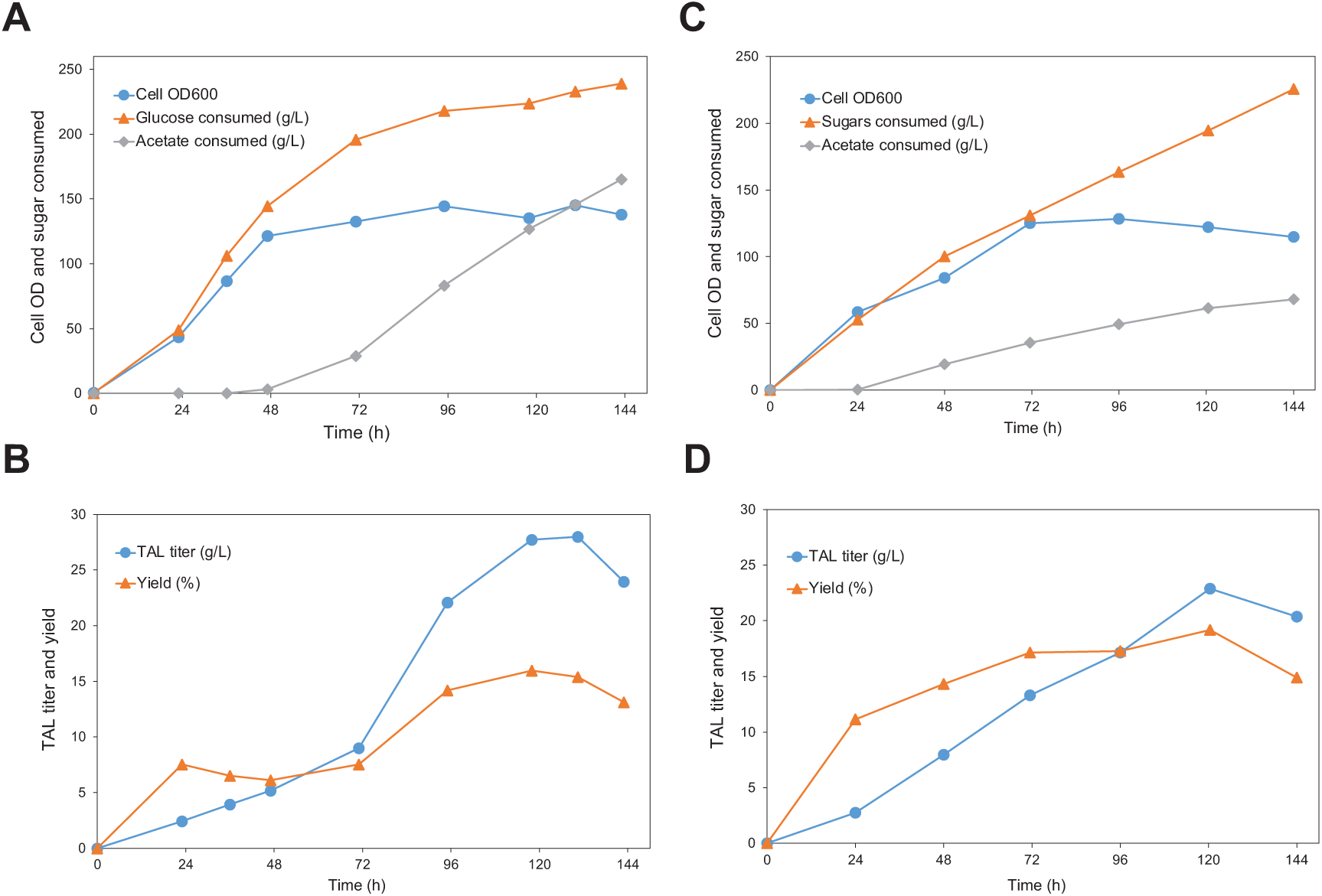
Fed-batch bioreactor fermentation of *R. toruloides*. A. The cell growth (OD_600_), total consumed glucose and acetate under glucose-based medium; B. The TAL titer and corresponding yield to its theoretical yield under glucose-based medium; C. The cell growth (OD_600_), total consumed glucose and acetate under oilcane juice-based medium; D. The TAL titer and corresponding yield to its theoretical yield under oilcane juice-based medium.

## 4. Conclusion

In this study, the codon-optimized 2-pyrone synthase (*GhPS*) was introduced into *R. toruloides* for TAL biosynthesis, and the resultant strain I12 produced ∼2 g/L TAL in culture tube. The dramatic improvement of TAL production by acetate addition suggests that acetate can not only serve as a substrate but also stimulate TAL production. It was found that *ACL1*, a citrate route enzyme, was a superior gene target to accumulate acetyl-CoA flux, and its overexpression improved TAL titer by 45% compared to I12 strain, which was 35% of the theoretical yield. The concurrent expression of *ACL1* and *ACC1* further improved TAL by 29% in culture tube, and the scale-up bioreactor fermentation achieved 28 g/L or 23 g/L TAL from glucose or low-cost oilcane juice with acetate spike, respectively. This work demonstrates that *R. toruloides* represents a promising microbial cell factory for production of polyketides and other acetyl-CoA-derived chemicals.

**Supplementary data to this article can be found online at xxx**.

## Author contributions

M.C. and H.Z. conceived the study and wrote the manuscript. M.C., V.T., C.S., and C.H. performed the experiments and analyzed the data. J.Q., A.O., and D.X contributed to bioreactor fermentation and discussion. All the authors proofread and agreed to publish.

## Acknowledgments

This work was supported by the U.S. Department of Energy award DE-SC0018420. The 1-L fed-batch fermentation experiments at UML was supported by the Acorn Innovation Grant. We thank Christopher Rao at the University of Illinois at Urbana-Champaign for the gifts of *R. toruloides* IFO0880 and plasmids pGI2 and pGI2_880_ACC. We also thank Vijay Singh at the University of Illinois at Urbana-Champaign for providing the oilcane juice and in-depth discussion on the setup of fermentation.

